# Evaluating Cell Identity from Transcription Profiles

**DOI:** 10.1101/250431

**Authors:** Nancy Mah, Katerina Taškova, Khadija El Amrani, Krithika Hariharan, Andreas Kurtz, Miguel A. Andrade-Navarro

## Abstract

Induced pluripotent stem cells (iPS) and direct lineage programming offer promising autologous and patient-specific sources of cells for personalized drug-testing and cell-based therapy. Before these engineered cells can be widely used, it is important to evaluate how well the engineered cell types resemble their intended target cell types. We have developed a method to generate CellScore, a cell identity score that can be used to evaluate the success of an engineered cell type in relation to both its initial and desired target cell type, which are used as references. Of 20 cell transitions tested, the most successful transitions were the iPS cells (CellScore > 0.9), while other transitions (e.g. induced hepatocytes or motor neurons) indicated incomplete transitions (CellScore < 0.5). In principle, the method can be applied to any engineered cell undergoing a cell transition, where transcription profiles are available for the reference cell types and the engineered cell type.

**Highlights:**

- A curated standard dataset of transcription profiles from normal cell types was created.
- CellScore evaluates the cell identity of engineered cell types, using the curated dataset.
- CellScore considers the initial and desired target cell type.
- CellScore identifies the most successfully engineered clones for further functional testing.

## Introduction

The discovery that a terminally differentiated cell type could be reverted to a pluripotent cell state capable of generating all possible cell types from the three germ cell layers (Takahashi and Yamanaka, 2006) has revolutionized the field of stem cell research and regenerative medicine. This concept of "reprogramming" one cell type to another has been further applied to lineage or direct reprogramming (Heinrich et al., 2015), in which cells are directly reprogrammed from one differentiated cell type to another, without first passing through a pluripotent state (Kim and Schöler, 2014). The development of these methods enable the production of patient-specific cells, which could be used for individualized drug testing and cell replacement therapy.

However, before these pluripotent stem cell-derived or other engineered cell types can be used for clinical applications, the safety and efficacy of the cells must be proven. First and foremost comes the patient’s safety with regard to oncogenicity and transplant rejection. For example, the process of creating iPS cells, including the reprogramming protocol and culture conditions, may introduce genomic aberrations (Lund et al., 2012), which could potentially render the iPS cells and their derivatives oncogenic. Indeed, the first phase 1 clinical trial using iPS cells was put on hold after gene mutations that were not present in the donor fibroblasts were found in the iPS cells used to treat age-related macular degeneration (Garber, 2015). Strict quality control of each patient-specific iPSC line for autologous transplantation also increases the cost and time for individualized treatment. In favor of a faster and more cost-effective way to generate clinical-grade cells, the suspended clinical trial aimed to resume (https://ipscell.com/2015/07/firstipscstop/) with the use of quality-controlled, banked, human leukocyte antigen (HLA)-typed iPS cells for allogeneic transplantation (Scudellari, 2016), which should reduce the risk of transplant rejection. In addition to successful safety trials, the cell-based treatment must demonstrate effectiveness. But even before the process of human clinical trials starts, engineered cell types that are destined for cell therapy must be sufficiently characterized. We propose that characterization of any kind of derived cell type that is intended for clinical use, can be partly accomplished by analyzing the gene expression profiles, in comparison to their starting donor cells and desired target cells. The same applies to their use for disease modelling and drug testing. Before extensive time and effort is spent on functional testing of engineered cell lines, transcription profiles could already be used as an initial screen to identify the most promising clones.

Current protocols to reprogram cells from one cell type to another typically involve forced expression of key factors, modulation of pathways with inhibitors and/or agonists, and changes in growth conditions. These protocols evolve from informed guesses tested by trial and error strategies, and even then, the molecular mechanisms behind cell transition are not completely understood. Parallel to the rise of reprogramming protocols, technological advancements allowing researchers to quantitate various cell properties in a high-throughput manner have created golden opportunities for the integration and analysis of "-omic" data. A large body of transcriptome data, i.e., quantitative measurements of the mRNA expression levels in a sample, has accumulated in the public domain, owing to high-throughput technologies such as microarray expression profiling or deep sequencing technology. Thanks to these technologies, there are millions of transcriptomic ’snapshots’ of cells, which can be used to define the state of a biological sample by computational means of determining the cell identity of engineered cell types. For example, Pluritest assesses the pluripotency of query cells by comparing the query with a standard expression matrix (of pluripotent stem cells and differentiated cells), using non-negative factorization models (Müller et al., 2011). In another method, CellNet quantifies the similarity of engineered cells and their target cell types, by extracting gene regulatory networks from transcriptome data of diverse cell types and further filtering the networks for regulatory interactions with ChIP-chip/-seq data (Cahan et al., 2014; Morris et al., 2014). Finally, KeyGenes uses a generalized linear regression model to extract classifier genes from a panel of transcriptional human fetal tissue, and then uses the "key genes" to classify query samples (Roost et al., 2015; Takasato et al., 2015).

In the present study, we propose a method to generate the CellScore. Unlike other computational methods, the CellScore method evaluates the cell identity of an experimentally derived cell type by taking into account both the initial donor cell type and the target cell type. First, we manually curated a set of reference transcriptome profiles from various normal tissues or cell types (Figure 1A). Then, we developed a method to assess cell identity of engineered cell types, using the expression profiles of the donor cell type and the desired target cell type as references (Figure 1B). Our method is targeted towards researchers following the gene expression changes of cells whereby the cells are expected to change their identity from a starting donor cell type towards a modified (e.g. reprogrammed or differentiated) cell type. If the desired target cell type has appropriate gene expression data, our method can be used to evaluate the “proximity” between an engineered cell type and its desired target cell, in terms of a CellScore. We can also evaluate whether the changes in gene expression during the identity change are leading to the desired state, and which genes contribute to this successful reprogramming. Finally, the method and accompanying reference dataset have been implemented as R packages that offer functions to calculate and visualize the cell scoring results for a given cell transition dataset in multiple ways including a single function call to obtain a detailed summary report in PDF format.

**Figure 1.**
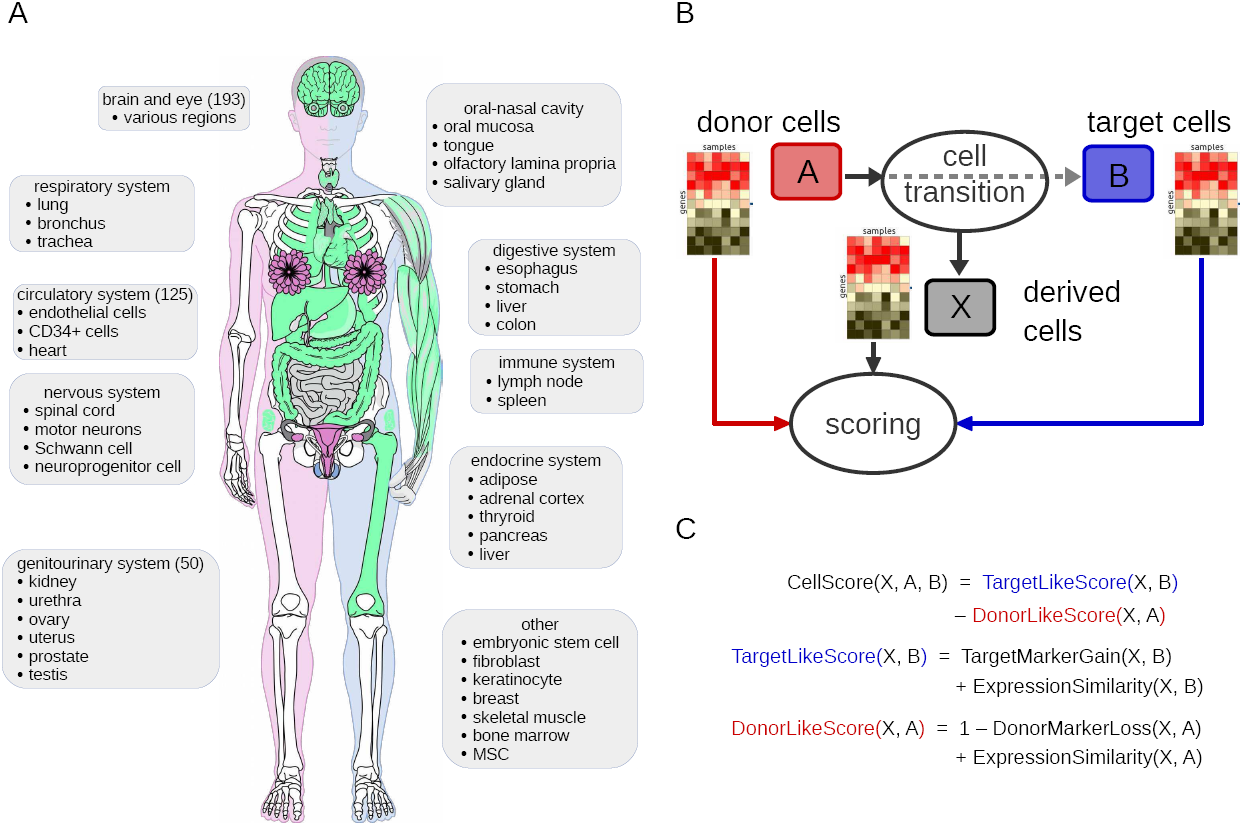
Reference cell types and the cell scoring strategy. A. The reference expression dataset consists of samples obtained from various tissues and cells, from public databases (see Supplementary Data S1 for a detailed list). This includes common (green) and sex-specific tissues or cells from male (blue) and female (pink) donors. Image of human body has been adapted from the on-line resource CellFinder (www.cellfinder.org). B. A derived cell type, X, is evaluated in comparison to its donor cell type A and target cell type B. C. The CellScore is the difference between the target-like and the donor-like score components. Each component score is calculated from the corresponding (either donor or target) marker gene expression and a measure of expression similarity, which in this study was chosen to be the cosine similarity.

## Results

### Transcriptome data across multiple studies defines a consistent expression space

To obtain a broad representation of expression space, studies involving cell transitions, as well as studies with normal cells were chosen for analysis. The reference dataset included over 100 standard cell types, consisting of either tissues, isolated cell types, or primary cell lines (Figure 1A, Supplementary Data S1). To demonstrate the utility of the CellScore, we compared over 300 test samples in transitions to over 10 target cell types of interest in regenerative medicine. Visualization of the reference expression dataset by principal component analysis (PCA; Figure 2) showed clusters of the same cell types, such as embryonic stem cells (navy blue; bottom left), fibroblasts (cornflower blue; left side) and brain or nervous tissues (pink and purple; right side), indicating that the normalization process was able to preserve sample characteristics, even though the samples originated from different studies.

**Figure 2.**
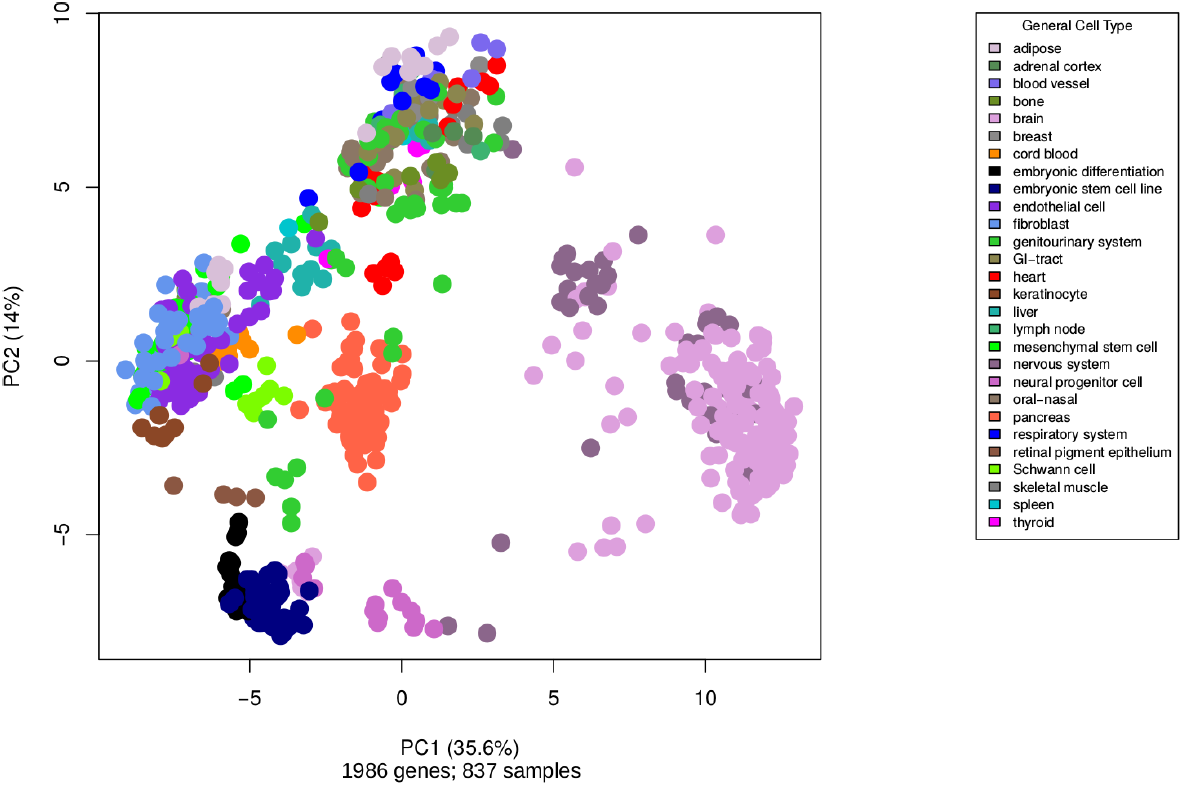
Principal component analysis plot of standard samples. Expression data from a wide variety of human tissues and cells were used as reference expression profiles for the cell scoring method.

### Similarity scores based on quantitative expression values

To further examine the similarity between donor, target, and engineered cell types, cells were sub-categorized according to their donor cell type to yield a cell subtype. From here on, we denote cell subtypes in the following format: derived cell type, “–”, donor cell type. Using the expression values, cosine similarity was calculated between the centroids of each subtype (Figure 3A). For the transitions to the ESC (embryonic stem cell) target cell type (i.e. iPS reprogramming), most of the iPS subtypes were highly similar to ESCs (Figure 3B). One notable exception were the "iPS-GSC*" cells, which were originally claimed to be iPS lines derived from spermatogonial cells (GSC), but were later shown to be more consistent with a fibroblast cell type (Ko et al., 2010). This is clearly visible in Figure 3B, where "iPS-GSC*" clusters with all other fibroblast cell types. A complete heatmap overview of the average cosine scores of all transitions is available in Supplementary Data S2.

**Figure 3.**
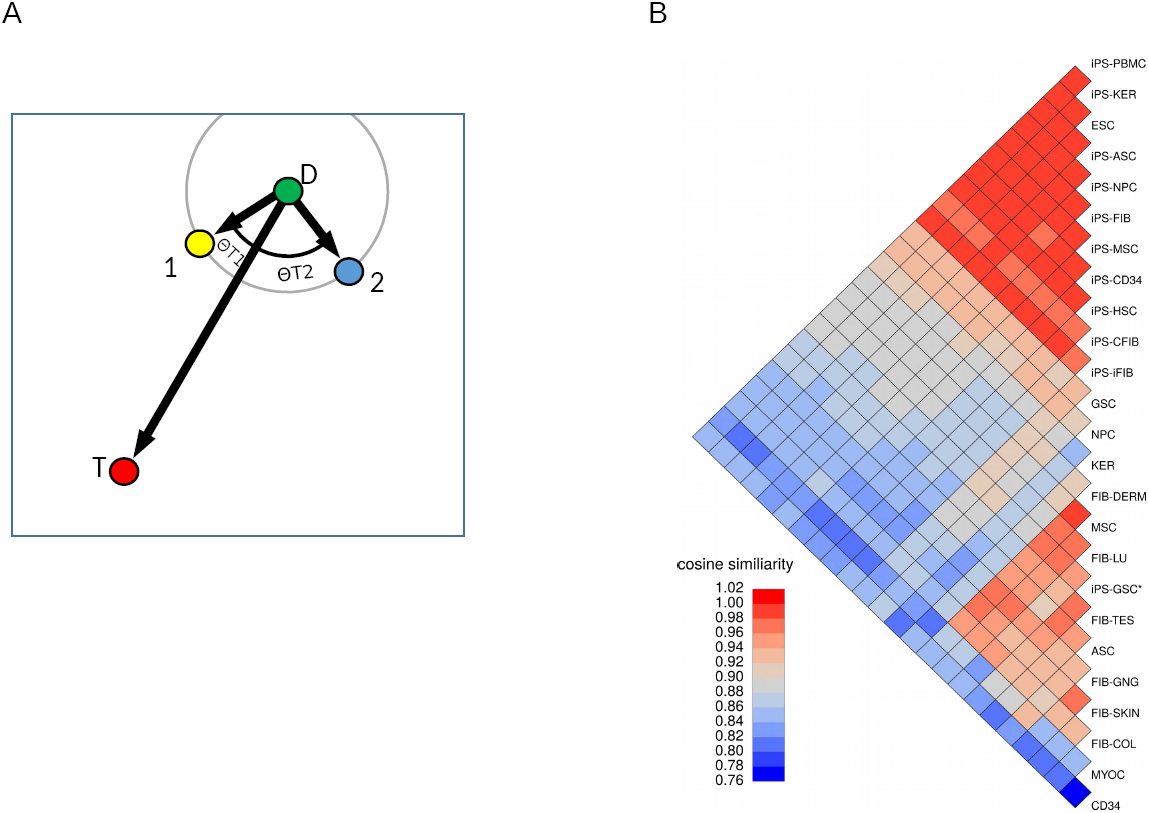
The use of cosine similarity as a cell scoring metric. A. For simplicity, the graph represents the gene expression of four samples projected in two-dimensional space. Point D is a donor cell type used for reprogramming. Point T is the desired cell type. Points 1 and 2 are engineered cell types, whose aim is point T. We would say that cells of point 1 are a more successful outcome of the transition from D to T, than cells of point 3 because cos(Θ_T1_) < cos(Θ_T2_). Note that such a comparison is different from using the Euclidean distance (grey circle around point D) to define similarity between samples. Our comparison takes into account the origin and desired endpoint of the cell transition. B. Heatmap of cosine similarity scores for all iPS reprogramming transitions. Average cosine similarity scores (across all cell type specific samples) are shown for various donor cells (mesenchymal stem cells, myocardium, CD34 positive cells from umbilical cord, keratinocytes, germ stem cells, neural progenitor cells and fibroblasts from dermis, lung, testes, gingiva, skin, colon), the target cell type (embryonic stem cells; ESC) and the iPS cells from each donor subgroup. The iPS cells have high cosine similarity scores compared to ESCs and cluster together at the top of the heatmap. Donor cell types originate from highly diverse tissues (bottom half of heatmap). The cell subtypes are formatted as: cell type, “-”, donor cell type or source tissue.

### Transition progression based on cell type markers

Aside from the expression data values, "present" calls from the microarray data analysis determine whether a gene was reliably detected or not. For each experimental cell subtype, we calculated the fraction of donor cell markers lost and target cell markers gain, to define an on/off score. Ideally, a successfully transitioned cell should lose all of its donor character and gain all of the target cell markers. For the most part, cells reprogrammed to iPS underwent a near complete loss of donor markers and gained at least 50% of the target markers (Figure 4A). Lower scoring transitions included those with transdifferentiation, highlighting the difficulty of these particular direct lineage reprogramming protocols to effect transition without first going to pluripotency. Examining the lists of donor and marker genes that are detected in the test cell types also provides valuable clues as to why a transition could not be carried to completion. As an example, functional enrichment in Reactome pathways was carried out on both partially reprogrammed iPS cells (piPS-FIB) and published iPS-FIB test cell types within the FIB (fibroblast) to ESC transition. The functional enrichment based only on these on/off genes demonstrated that certain Reactome categories, such as "Signaling by NODAL" or "Transcriptional regulation of pluripotent stem cells" were clearly lacking expression in the partially reprogrammed cells compared to ESCs, whereas the iPS-FIB cells showed an enrichment profile very similar to that of ESCs (Figure 4B).

**Figure 4.**
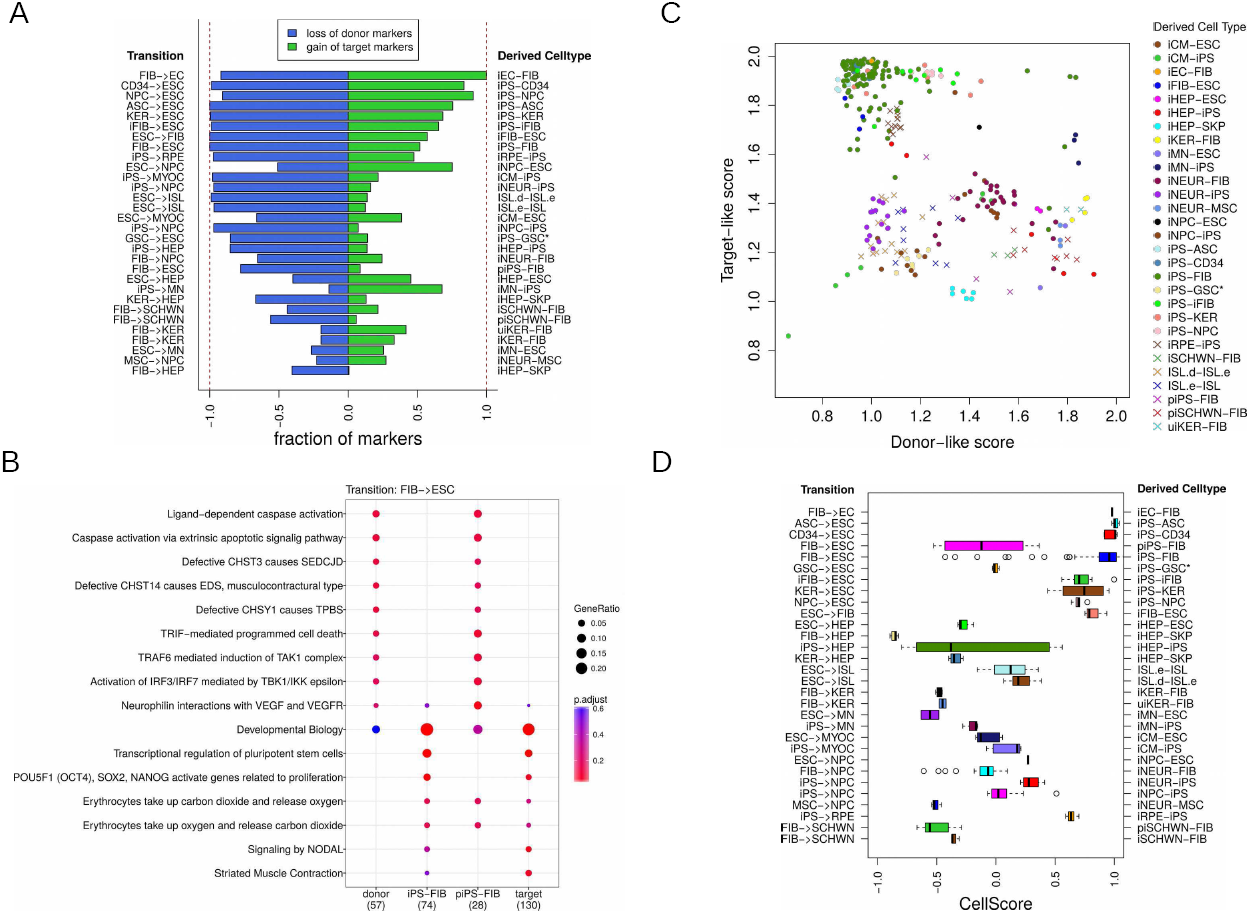
CellScore metrics for selected transitions. A. Barplot of on/off scores for selected transitions. A complete loss of donor markers (blue) and a gain of target cell markers (green) indicates the most successful transition between two cell types. Shown are all derived cell types for transitions targeting embryonic stem cells (ESC), fibroblasts (FIB), hepatocytes (HEP), keratinocytes (KER), myocardium (MYOC), retinal pigmented epithelium (RPE), motor neurons (MN), Schwann cells (SCHWN), endothelial cells (EC) and neuroprogenitor cells (NPC). The experimentally derived cell types are formatted as: derived cell type, “-”, donor cell type. Bars indicate total fractions of lost (depicted as a negative value for donor markers) or gained markers (shown as a positive value) for all samples of one derived cell type. B. Functional enrichment of on/off markers for the transition from donor fibroblasts to target embryonic stem cells. Functional enrichment for two types of test cells (partially induced iPS cells and iPS cells, both from fibroblasts) was carried out with the Reactome pathway set. piPS-FIB cells share an enrichment profile most similar to the donor fibroblast cells. iPS-FIB cells are more similar to the ESCs. Notably, of the 28 target marker genes in the category ‘Developmental Biology’, the iPS-FIB cells express 18 of these, whereas the piPS cells only express four of the target marker genes. Dot size indicates the proportion of genes present in each cell type per universe set of genes, which was defined to be the set of all present genes in the standard data matrix. Enrichment p-values (low to high p-values represented by a gradient of red to blue, respectively) for each Reactome category were corrected for false discovery rate (Benjamini and Hochberg method). C. Scatterplot of donor-like and target-like scores is shown for 30 experimentally derived cell types. Each point represents an individual cell line. The most successful transitions have low donor-like scores and high target-like scores. D. Distributions of CellScore values for the same 30 cell transitions are shown as boxplots. Positive CellScore indicates that the derived cell type is more similar to the desired target, whereas negative CellScore indicates a higher similarity to the donor cell type. The experimentally derived cell types are formatted as: derived cell type, “-”, donor cell type. Abbreviations: induced pluripotent stem cell (iPS), embryonic stem cell (ESC), mesenchymal stem cell (MSC), induced cardiomyocyte (iCM), induced fibroblast (iFIB), iHEP (induced hepatocyte), skin-derived precursor (SKP), induced keratinocyte (iKER), induced motor neuron (iMN), induced neuron (iNEUR), induced neural progenitor cell (iNPC), adipose stem cell (ASC), CD34+ cord blood cell (CD34), germ stem cell (GSC), induced retinal pigmented epithelium (iRPE), induced Schwann cells (iSCHWN), partially induced Schwann cells (piSCHWN), expanded islet cells (ISL.e), re-differentiated expanded islet cells (ISL.d), fresh islet cells (ISL), partially induced iPS cell (piPS), undifferentiated induced keratinocyte (uiKER).

### Application of the cell scoring method

The CellScore is composed of target‐ and the donor-like scores (Figure 1C), which are obtained from the cosine similarity and on/off (marker gain/loss) score. The CellScore values, including the values of the specific score components, are shown for 30 selected transitions in Figure 5. The engineered cells with the highest similarity to their desired target cell types are concentrated in the upper-left corner of Figure 5A, with high target-like scores (>1.5) and low donor-like scores (< 1.5). The CellScore is simply the difference between the target‐ and donor-like scores, as shown in Figure 5B. The CellScore ranges between −1.2 and 1.2. A highly positive CellScore indicates that the experimental cell is very similar to its target cell type, while a highly negative score indicates that the experimental cell type has not successfully transitioned and has remained more donor-like. To visualize the CellScore and the metrics that contribute to the CellScore of an engineered cell type, a report can be generated with diagnostic plots as in Figure 5. Reports for all cell transitions in the test dataset were generated and are available as Supplementary Data S3, and the complete CellScore values and marker gene lists are available as Supplementary Data S4. Though many transitions were assessed in the test dataset, we demonstrate the utility of the approach to score experimental cell types in the following selected examples:

**Figure 5.**
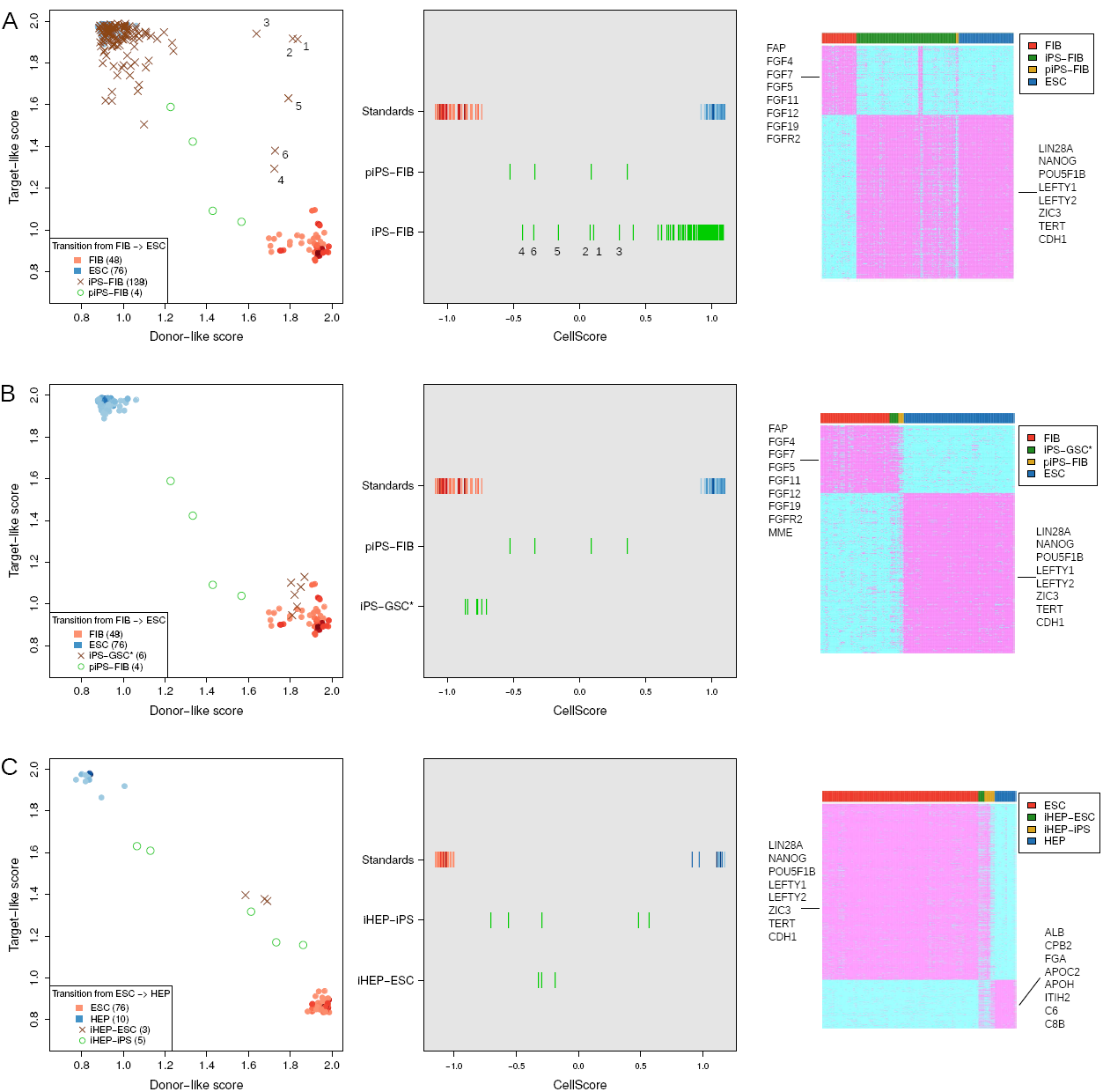
CellScore case studies. Each row of figures represents a case study, and each column shows diagnostic plots for the case study. The first column on the left shows scatterplots of the donor-like and target-like scores, with donor standards and target cell standards in red and blue colours, respectively. The colours of the standard samples are shaded darker, according to the number of overlapping points. The middle column shows rug plots of CellScore values, comparing CellScore values of the derived cell types with the CellScore values of the donor and target standard samples. Finally, the column on the right are heatmaps depicting the marker gene expression, where marker gene detection is indicated by magenta (present) and cyan colours (absent), respectively. Samples in columns are grouped by donor (red bar), test (green/yellow bar) or target (blue bar). Short lists of some of the genes in each marker category are shown on the side of each heatmap. A. FIB to ESC transition. iPS cells derived from fibroblasts, across all studies, are shown. Partially induced iPS cells from FIB (piPS-FIB) are included for comparison. Numbers indicate outlier samples from study GSE27206. B. Special case of iPS-GSC* cells in the context of FIB to ESC transition. C. ESC to HEP transition.

Case 1: FIB to ESC. Within the context of the fibroblast-derived iPS cells, most iPS cells (iPS-FIB) indeed have target-like scores and low donor scores, leading to high CellScore values (Figure5A). Partially reprogrammed iPS cells (piPS-FIB) show gradual transition to ESC, and have CellScore < 0.5, indicating their incomplete transition. Notably, there are six outlier iPS-FIB samples from the same study (GSE27206) (Corti et al., 2012) with high donor-like scores on the right side of Figure 5A, first panel. Their CellScore values range from −0.43 to 0.30, and display both FIB marker and ESC marker expression (Figure 5A, middle and right panel). Two of the iPS lines originate from spinal muscular atrophy patients (#1,2 in Figure 5A), one is a wild-type parent of the patients (#3) and the remaining iPS lines (#4,5,6) were previously established lines (iPS 19-9; hPSCreg identifier WAi001-A) from the Thomson laboratory (Yu et al., 2009). We also have calculated the CellScore for the iPS 19-9 lines in Supplementary Data S3 (see GSE15176). All iPS 19-9 lines from the original study (Yu et al., 2009) have high CellScore values (0.96-1.05) and are virtually indistinguishable from ESC standards. Together these results indicate a potential systematic problem in the iPS cells profiled in GSE27206. It should be noted these outlier cell lines were in turn used as starting material for the production of patient-derived motor neurons; therefore these cells could be considered as transdifferentiation intermediates rather than fully reprogrammed iPS lines. Overall, of the 138 iPS-FIB samples assessed by CellScore, 72% had values greater than 0.9, which indicates high transition to an ESC-like state. The iPS-FIB cell lines with lower CellScore have not silenced the expression HOX family body patterning genes, which are highly expressed in the donor fibroblasts (Figure 6). Expression of a higher number of HOX genes in the iPS cell lines tends to decrease the CellScore and indicates a less successful transition.

**Figure 6.**
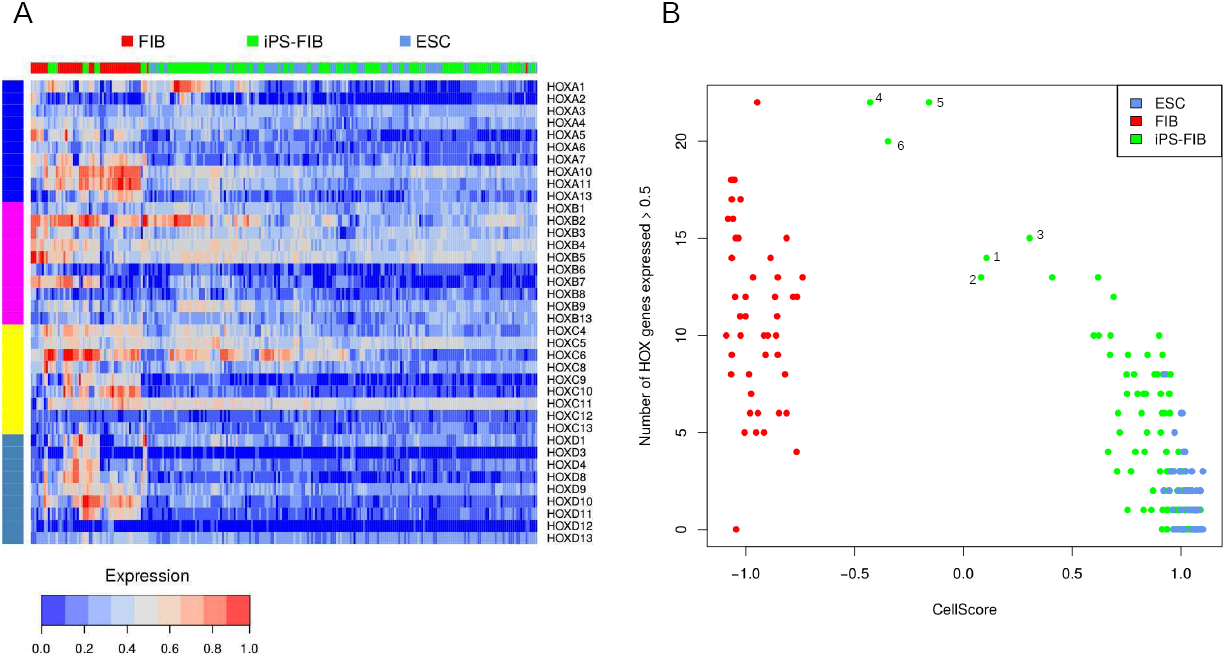
HOX gene expression in iPS-FIB cells. A. Heatmap of normalized expression in HOX gene families for fibroblasts (FIB), induced pluripotent stem cells reprogrammed from fibroblasts (iPS-FIB), and embryonic stem cells (ESC). B. Scatterplot of CellScore values of cell types according to the number of HOX genes with expression greater than 0.5. Numbered points on the scatterplot indicate the outlier samples from study GSE27206 as in Figure 5A.

Case 2: GSC to ESC. Interestingly, in one well-known controversial case of iPS cells, which were supposed to have been derived from spermatogonial cells (iPS-GSC*; GSE11350) (Conrad et al., 2008), the derived cells had a CellScore of about zero in the transition from spermatogonial germ cells to ESC (Figure 5B). However, when these iPS-GSC* cells were scored in the FIB to ESC transition, the CellScore values were very close to values of the fibroblast cell line standards (Figure 5B). These cells have been shown to be most likely testicular fibroblasts (Ko et al., 2010) and not iPS cells.

Case 3: ESC to HEP. We analyzed three studies of converting pluripotent stem cells to hepatocyte-like cells (GSE62962, GSE14897, GSE62547). Each study used a different method to produce the induced hepatocytes (iHEP) and CellScore provides an objective way to compare their outcomes. The most successful conversion of PSC to HEP came from GSE62962, where the authors used a micropatterned co-culture platform with stromal fibroblasts to differentiate the cells to iHEP (Berger et al., 2015). Though the CellScore for these two lines was only 0.57 and 0.48, the lines clearly had moderate target-like scores and low donor-like scores (Figure 5C). In the microarray expression data, the iHEPs from this study expressed high levels of albumin, like the primary hepatocytes that were used as standards. However, a high level of alphafetoprotein gene in the iHEP cells shows that they were still immature hepatocytes and had not completely converted to the desired target cell type, primary hepatocytes. Nevertheless, the authors demonstrated that their iHEP cells had hepatocyte morphology and polarity, as well as drug metabolizing ability (Berger et al., 2015).

### Performance of the method

To evaluate the performance of the CellScore method, we curated an independent dataset of primary cells, cell lines or tissues from 25 studies, which were not previously used as standards for CellScore. ESC, HEP, and RPE (retinal pigmented epithelium) were then defined as true positives in three transitions (FIB → ESC, ESC → HEP, ESC → RPE), respectively, to calculate Receiver Operating Characteristic (ROC) curves (Figure 7). Other cell types within these 25 studies were used as true negatives, with the exception of iPS cells, which were too similar to ESCs. Within a transition, all samples that were not true positives were used as true negatives. All three transitions yielded Area Under the Curve (AUC) values greater than 0.97, which indicates excellent performance in classifying the tested cell types. Both ESC and RPE cells were classified with very high sensitivity (0.95) and high specificity (false positive rate – FPR of 0.05), whereas the HEP classification had a higher FPR (0.14) at 0.95 sensitivity. This higher FPR was partly driven by chemically treated hepatocyte samples, which were counted as true negatives, but in fact were not significantly influenced by small molecule treatment.

**Figure 7.**
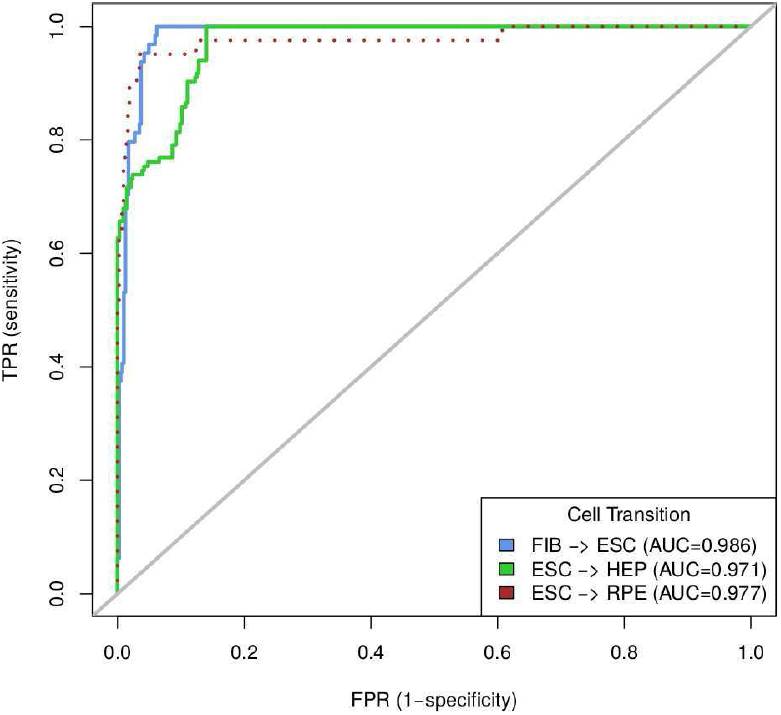
Receiver Operating Characteristic (ROC) curve for CellScore. ROC curves were generated for three transitions: FIB → ESC (blue), ESC → HEP (green), and ESC → RPE (brown). Grey line indicates random performance. True positives for each transition are the target cell type of each transition. True negatives for each transition are the donor cell type and a variety of cell types, including modified ESC, embryonic carcinoma, induced hepatocytes, treated RPE, Wharton’s jelly stem cells and cancer cell types (brain, bladder, colon, lung, prostate).

## Discussion

In the present study, we propose a new approach, called CellScore, to score experimental cell types undergoing a transition in cell identity. This is a particularly emerging current issue as it becomes more routine to engineer an essential cell type on-demand for disease modelling or patient-specific drug-testing. The CellScore alone is not intended to wholly replace other cellular and molecular techniques to confirm the desired cell function, but CellScore results can already provide some information about the success of engineered cell transition.

We applied the CellScore method to over 20 transitions in cell identity from data available from one particular microarray platform, including over 1500 samples in 86 studies. Some transitions were very popular and had much data from multiple studies (e.g. FIB →ESC, PSC → HEP, PSC → MYOC) while others, such as those deriving cells of the nervous system, only had data from a single study. In three transitions, for which additional public data was available from the same platform, we were able to show that the CellScore method was able to classify true positives with high specificity and low false positive rates. Overall, and probably not surprising, the highest scoring engineered lines were the iPS lines and other transitions involving cultured cell lines as standards (e.g. ESC → FIB, FIB → EC). Other transitions did not reach CellScore values above 0.5. There could be many explanations for these incomplete transitions. First, there could be experimental inconsistencies in the generation or maintenance of the derived lines. The meta-analysis of the test iPS samples detected clear outliers: the known problematic cell lines in the case 3 from above (Conrad et al., 2008) and the unusual iPS lines of case 1 from above (Corti et al., 2012), in which all six iPS lines were affected with low CellScore values, including three bona fide iPS lines from another laboratory. Second, differentiation protocols to produce specialized cell types are constantly being improved, and engineered cells in the current analysis do not reflect cells generated from the most recent advances in vitro cell differentiation. Finally, there might not be suitable target cells available for the CellScore method. Engineered cells are typically grown in culture, and these conditions cannot possibly mimic the cell types in their native in vivo environments. Many engineered cell types could be considered more fetal-like, rather than terminally differentiated, and due to ethical and practical considerations, there is a lack of sufficient numbers of human fetal tissue samples that could serve as standards. Therefore, the standard target cell type used to calculate the CellScore is a proxy, and in itself represents an ideal and unattainable goal. For example, in the PSC → HEP transition, primary hepatocytes were used as the target; however, the induced hepatocytes were still immature compared to primary hepatocytes. In spite of their immaturity, these cells already demonstrated sufficient functionality in terms of metabolic activity and liver marker expression, such that they could be used in some in vitro toxicity assays (Berger et al., 2015). It may not even be necessary to derive mature hepatocytes, depending on its downstream use. Immature hepatocytes could be used as a form of cell replacement therapy by injecting them into a liver, where the immature hepatocytes could fully differentiate into hepatocytes in situ (Si-Tayeb et al., 2010). In such cases, the CellScore could still be useful as a guide to choosing the most differentiated iHEPs and compare outcomes between distinctly scoring cells.

Beyond the CellScore as a single number to quantify a cell transition, the CellScore reports give additional information showing the contribution of target-like and donor-like metrics to the CellScore and diagnostic heatmaps of the markers expressed by standards and experimentally derived cells. As exemplified by the case of partially induced iPS cells, the on/off marker lists can be extracted and further analyzed to highlight genes responsible for an incomplete transition, which could be in fact driven by donor cell type-specific transcription networks (Nefzger et al., 2017) Functional enrichment analysis could highlight pathways or processes that may be suitable input for small molecule induction/repression prediction, in an effort to improve differentiation protocols (Siatkowski et al., 2013).

Other computational methods to evaluate cell identity exist, but mainly these are restricted in that they perform specific functions and rely on specific technologies. Pluritest focuses on the scoring of human pluripotent stem cells and is restricted to using datasets from dedicated microarray platforms, as a means for standard testing on a web-based platform behind a login-wall (Müller et al., 2011). Keygenes is a specialized resource for scoring RNA-seq profiles to a fixed panel of human embryonic tissues, with freely available software (Roost et al., 2015). On the other hand, CellNet is applicable to microarray data of different platforms, provided that at least 60 microarrays are available for each tissue type. CellNet uses cell or tissue-specific gene regulatory networks to classify engineered cell types and its software is freely available (Cahan et al., 2014). None of these methods explicitly takes into consideration the donor cell type when evaluating the cell identity. We propose the CellScore method as a freely available method to evaluate any kind of cell identity transition, which could in principle score any type of cell transition for which donor cell and target cell data are available, regardless of the platform. The current study has been limited to data from one platform as a proof-of-principle, but in the future, we aim to leverage the data from additional microarray platforms, as well as RNA-seq datasets, for a comprehensive expression database (Mah et al., 2017) to evaluate cell identity.

## Experimental Procedures

### Data selection and processing

Transcriptome data was collected from published studies that involved changes in cell identity, i.e. reprogramming somatic cells to iPS, directed differentiation of pluripotent cells, or direct lineage reprogramming. Donor cells were defined to be the starting cell types, whereas target cells were the desired endpoint cell types. Engineered cell types were defined to be those undergoing the cell transition from the donor to the target cell type. Further studies were added as necessary to complement the number of transcription profiles from standard (donor or target) cell types. This data collection contains pluripotent stem cells, adult stem cells and differentiated cells, which serve as standard cell types to which the engineered cells are compared. An overview of the reference samples in the data collection is shown in Figure 1A and a complete list of all samples is given in Supplementary Data S1. Samples were manually annotated for their tissue origin and cell types using ontology terms (Malone et al., 2010; Stachelscheid et al., 2014). Raw microarray data were obtained from the public repositories Gene Expression Omnibus and ArrayExpress (Barrett et al., 2009; Rustici et al., 2013). Microarray data was normalized using the YuGene transform, which allows comparisons between experiments and is not dependent on the data distribution (Lê Cao et al., 2014). For present/absent calls, MAS5.0 detection p-values were calculated using the "affy" Bioconductor package for Affymetrix 3’IVT arrays (Gautier et al., 2004). Probesets in each sample were considered to be "present" if the detection p-value was less than 0.05. Annotation of probesets to genes was obtained from the BioC annotation package "hgu133plus2.db" (Carlson M, 2016). In the case that multiple probesets mapped to one gene, the probeset with the highest median across all samples was chosen to represent the expression for that gene.

### Cell scoring method

Cell scores for a given cell transition starting from a specific cell type (defined by the donor cell type) towards a different cell type (defined by the target cell type) were calculated using two independent metrics, the on/off score and the cosine similarity score (Figure 3A). Both metrics are composite, calculated once with the respect to the donor and once with the respect the target cell type (Figures 1C). The first metric was based on the so-called on/off genes, i.e. cell type specific genes that were uniquely expressed (detected as "present") in either the donor cell type (donor markers) or the target cell type (target markers) of a particular cell transition. Then the on/off scores for a query sample *i* was defined by the fraction of lost donor markers (*f_D,i_*) and the fraction of gained of target markes (*f_T,i_*), such that:

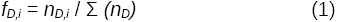

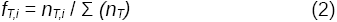

where *n_D,i_* and *n_T,i_* are the numbers of donor or target markers present in the sample *i*, and *n_D_* and *n_T_* are the numbers of donor and target markers, respectively, present in the standard cell types of a particular cell transition.

In the second metric, cosine similarity of transcription profiles was calculated between all target, donor and experimental cell types, based on a subset of the most variable genes from the standard cell types in the normalized expression matrix. By default, the most variable genes were defined to be those genes whose standard cell type group medians were in the top 10% of the expression interquartile range. Cosine similarity was calculated between each query sample and the mean centroid of each standard cell type. To get an intuitive impression of the cell identity of a query sample *i*, the donor-like score (*s*_D,_*_i_*) and the target-like score (*s_T,i,_*) were calculated as follows:

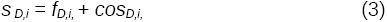

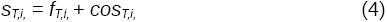

where *cos_D,i_* is the cosine similarity between the query sample *i* and the centroid of the donor cells, and *cos_T,i_* is the cosine similarity between the query sample *i* and the centroid of the target cells of a given cell transition. As both “on/off” score and the cosine similarity have values in the range of [0,1], the derived (donor-like and target-like) scores can have values in the range [0,2]. A successful cell transition will have samples with very high donor-like scores and very low target-like scores.

Finally, the CellScore of a query sample *i* for a particular cell transition was defined to be:

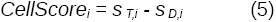

The CellScore values range between −2 and 2, with negative values indicating that the query sample is more donor-like (the cell transition was not successful), and positive values indicating that it is more like the target cell type (meaning a potentially successful cell transition). The cell scoring method and pre-processed reference and test data presented in this study is publicly available at https://github.com/nmah/CellScore.

### Functional enrichment and data visualization

The Bioconductor package "ReactomePA" was used to determine the enrichment of marker genes in Reactome pathways, using the entire list of genes in the dataset as the background (universe) set of genes (Yu and He, 2016). Enrichment results were plotted using the Bioconductor package "clusterProfiler" (Yu et al., 2012). Receiver Operating Characteristic curves were plotted by the R-package "pROC" (Robin et al., 2011).

### Generation of the CellScore Report

Users can produce a report summarizing the CellScore values for engineered cells in a particular cell transition. The report contains four diagnostic figures: 1) a scatterplot of the target‐ and donor-like scores of the engineered cells; 2) a density plot of the CellScore values for donor, target and engineered cell types; 3) a rug plot showing a detailed view of the engineered CellScore values in relation to the target cell CellScore values; 4) a heatmap of the on/off marker expression in the donor, target and derived cell types. Figures can also be plotted individually using the functions in the CellScore R-package. All calculations and plots were carried out using R-statistical software version 3.0 or greater (R Core Team, 2017).

## Author Contributions

NM, conception and design, collection and curation of data, software coding and testing, data analysis and interpretation, manuscript writing, and final approval of manuscript. KT, software coding and testing, software package development, manuscript writing and final approval of manuscript. KEA, software testing, manuscript writing and final approval of manuscript. KH, manuscript writing and final approval of manuscript. AK, manuscript writing and final approval of manuscript. MA, conception and design, manuscript writing and final approval of manuscript.

## Acknowledgements

We acknowledge the generous support from the German Stem Cell Network for travel awards to NM for stem cell-related meetings. We thank Prof. Petra Reinke for her continuing support. This work is supported by the European Commission grant no. 334502 (hPSCreg) and the Innovative Medicines Initiative Grant for the European Bank for induced Pluripotent Stem Cells (EBiSC).

## Tables

None.

## Supplemental Information

Supplementary Data S1. Spreadsheet of all samples used for reference and derived cell types.

Supplementary Data S2. Cosine similarity score heatmap for engineered cells and all standard cell types.

Supplementary Data S3. CellScore reports for all cell transitions and studies.

Supplementary Data S4. CellScore values and OnOff marker tables.

